# Compositional Flexibility of the ER-Mitochondria Encounter Structure

**DOI:** 10.1101/2024.11.26.625358

**Authors:** Christian Covill-Cooke, Takashi Hirashima, Shin Kawano, Joe Ganellin, Andrew Moody, Sabine N.S. van Schie, Arun T. John Peter, Chika Saito, Toshiya Endo, Benoît Kornmann

## Abstract

Yeast mitochondria receive the majority of their lipids from the endoplasmic reticulum (ER) via the heterotetrameric ERMES lipid transport complex. This complex is thought to establish a lipid transporting tube of fixed composition spanning the space between both organelles. Intriguingly, however, some of the lipid-transporting components of the complex can be replaced by an artificial ER-mitochondria tether without lipid transport activity, indicating that ERMES subunits are not all of equal importance for lipid transport. Here, we propose a model whereby lipid transfer by the ERMES complex can occur with various sub-ensembles of ERMES, and minimally with only one of the four members, namely Mmm1. Our results imply flexibility in the composition of the ERMES complex, which might help it accommodate various interorganelle distances.

## Introduction

The transport of lipids from their site of synthesis – predominantly the ER – to the mitochondrial membranes is essential for the existence of mitochondria and, therefore, eukaryotic life (Tatsuta and Langer, 2017; Tamura et al., 2020). In yeast, the transport of lipids from the ER to mitochondria is thought to occur predominantly through the ERMES complex, a heterotetrameric complex of Mmm1, Mdm12, Mdm34 and Mdm10 (Kornmann et al., 2009; Kornmann, 2020; John Peter et al., 2022). The ability of ERMES to traffic lipids is bestowed by lipid-transfer Synaptotagmin-like Mitochondrial-lipid-binding Protein (SMP) domains found in three of the members of the complex (Mmm1, Mdm12 and Mdm34). SMP domains form hydrophobic pockets that can shelter lipid from the aqueous cytosol, thus catalysing interorganelle lipid exchange (AhYoung et al., 2015; Kopec et al., 2010; Kawano et al., 2018; AhYoung et al., 2017; Jeong et al., 2017, 2016).

Important questions remain unaddressed regarding the biology of the ERMES complex. First, how does it transfer lipids at the mechanistic level? A crystal structure of a Mmm1-Mdm12 heterotetramer (homodimer of heterodimers) has been described (Jeong et al., 2017). Given the size of this heterotetramer, it has been proposed to shuttle lipids between both membranes using flexible linkers in Mmm1 and/or Mdm34. By contrast, recent cryotomography and structural prediction fitting (Wozny et al., 2023) indicates that the SMP domain of Mmm1 (anchored in the ER membrane) connects to that of Mdm12, which itself binds to Mdm34 to bridge over to Mdm10, embedded in the outer mitochondrial membrane (OMM). In this model, a linear and compositionally rigid Mmm1-Mdm12-Mdm34-Mdm10 heterotetramer constitutes a hydrophobic conduit for lipid transfer between the two membranes (Figure 1A). Yet, because the folded domains of Mmm1, Mdm12 and Mdm34 are similar in shape and size, at the resolution of the tomograms, one cannot exclude the existence of a variety of complexes with different subunit compositions, like Mmm1-Mdm12-Mdm12-Mdm10, or Mmm1-Mdm34-Mdm34-Mdm10, for instance.

**Figure 1:**
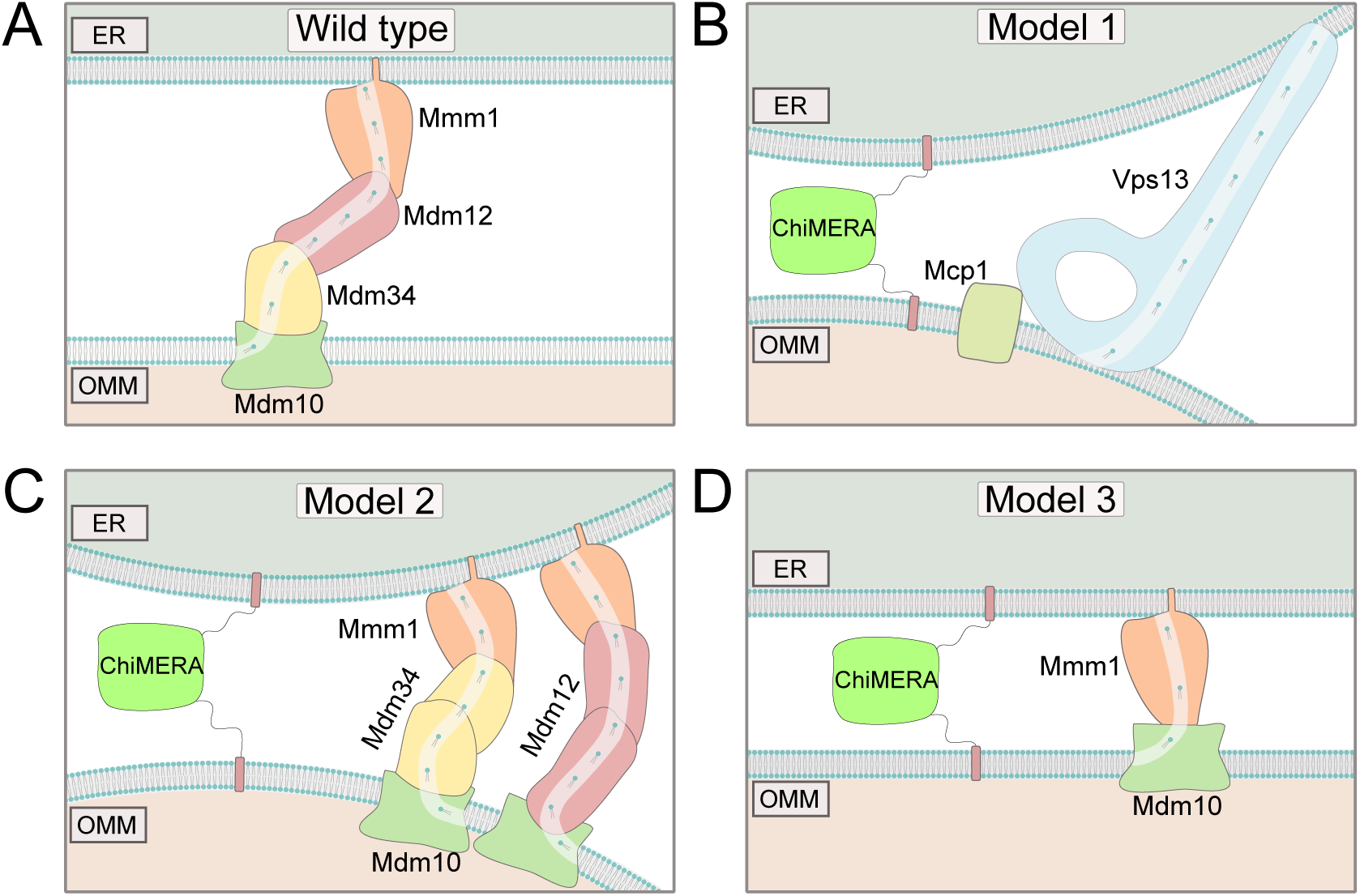
Models for ChiMERA rescue of ERMES depletion. (A) Model of the heterotetrameric ERMES complex. (B) Model 1: The lipid transporter Vps13 is recruited to artificial ER-mitochondria contacts. (C) Model 2: Mdm12 and Mdm34 are essential for tethering but redundant in lipid transport function. (D) Mmm1 and Mdm10 can support lipid transfer providing sufficient external tethering force.

Deletion of any of its members leads to the disappearance of ERMES structures, slow growth and mitochondrial malfunction, but does not entirely prevent cell growth in budding yeast (Kornmann et al., 2009; Kornmann, 2020; Burgess et al., 1994; Boldogh et al., 2003; Sogo and Yaffe, 1994; Berger et al., 1997). Indeed, ERMES mutant cells harbour mitochondria that are dysfunctional but nevertheless bounded by lipid-containing membranes, indicating that ERMES is not strictly necessary to provide lipids to mitochondria. Another complex involving the lipid transport protein Vps13 and its mitochondrial adaptor Mcp1 has been shown to act in partial redundancy with ERMES, to support lipid transport between the endomembrane system and mitochondria, and promote cell growth (John Peter et al., 2017; Lang et al., 2015; John Peter et al., 2022; Park et al., 2016). Increasing lipid transport through this pathway, for instance by overexpressing Mcp1 and forcing Vps13 recruitment to mitochondria, rescues the phenotypes associated with the lack of ERMES (John Peter et al., 2017; Tan et al., 2013).

The ERMES complex was first identified as an ER-mitochondria tethering complex because its absence can be suppressed by the expression of ChiMERA – a synthetic fusion protein that artificially tethers the ER to the mitochondria. More specifically, ChiMERA expression rescues the growth defect of *mdm12* and *mdm34* single mutants although it fails to rescue that of *mmm1* and *mdm10* mutants (Kornmann et al., 2009). Crucially, ChiMERA lacks lipid transport activity, at odds with the idea that ERMES is a lipid transport catalyst (Kornmann et al., 2009). This discrepancy can be explained by any of three potential models: 1) ChiMERA-induced tethering might exert its rescuing effect by allowing Vps13 recruitment to artificial ER-mitochondria contact, similar to Mcp1 overexpression, leading to rescue of ERMES mutants (Figure 1B); 2) While the presence of both Mdm12 and Mdm34 might be necessary for ERMES tethering function, both proteins might be redundant for lipid transfer in the presence of ChiMERA, as could be expected if Mmm1-Mdm12-Mdm12-Mdm10, or Mmm1-Mdm34-Mdm34-Mdm10 complexes of lower affinities existed (Figure 1C); 3) Mmm1 might be able to serve as the sole lipid transport protein (LTP) at ER-mitochondria contact sites established by ChiMERA-mediated tethering (Figure 1D). Distinguishing between these three models has important implications for the mechanism of ERMES-mediated lipid transport.

## Results

### Vps13 localizes at ER-mitochondria contacts but is dispensable for ChiMERA-mediated ERMES rescue

Model 1 states that ChiMERA expression recruits Vps13 to artificial ER-mitochondria contacts (Figure 2A). Although we have previously suggested that Vps13 rescued ERMES by connecting mitochondria to vacuoles, (John Peter et al., 2017), we found that Vps13’s vacuolar adaptor, Ypt35, is not necessary for Vps13-mediated ERMES rescue (Supp. Figure 1A) (Bean et al., 2018; Michel et al., 2017) nor for its recruitment to vCLAMPs upon Vps39 overexpression (Supp. Figure 1B), indicating that Vps13 likely binds promiscuously to membrane apposition sites upon Mcp1-mediated recruitment. Without Vps39 overexpression, we observed that Vps13^GFP was not homogeneously localized on the mitochondrial surface. Instead, Vps13^GFP appeared to form distinct bright puncta (Supp. Figure 1C). These bright foci of Vps13 colocalized very often with ERMES, significantly exceeding the frequency expected by chance (Figure 2B-C), providing a novel localization for Vps13 at ER-mitochondria contact sites.

**Figure 2:**
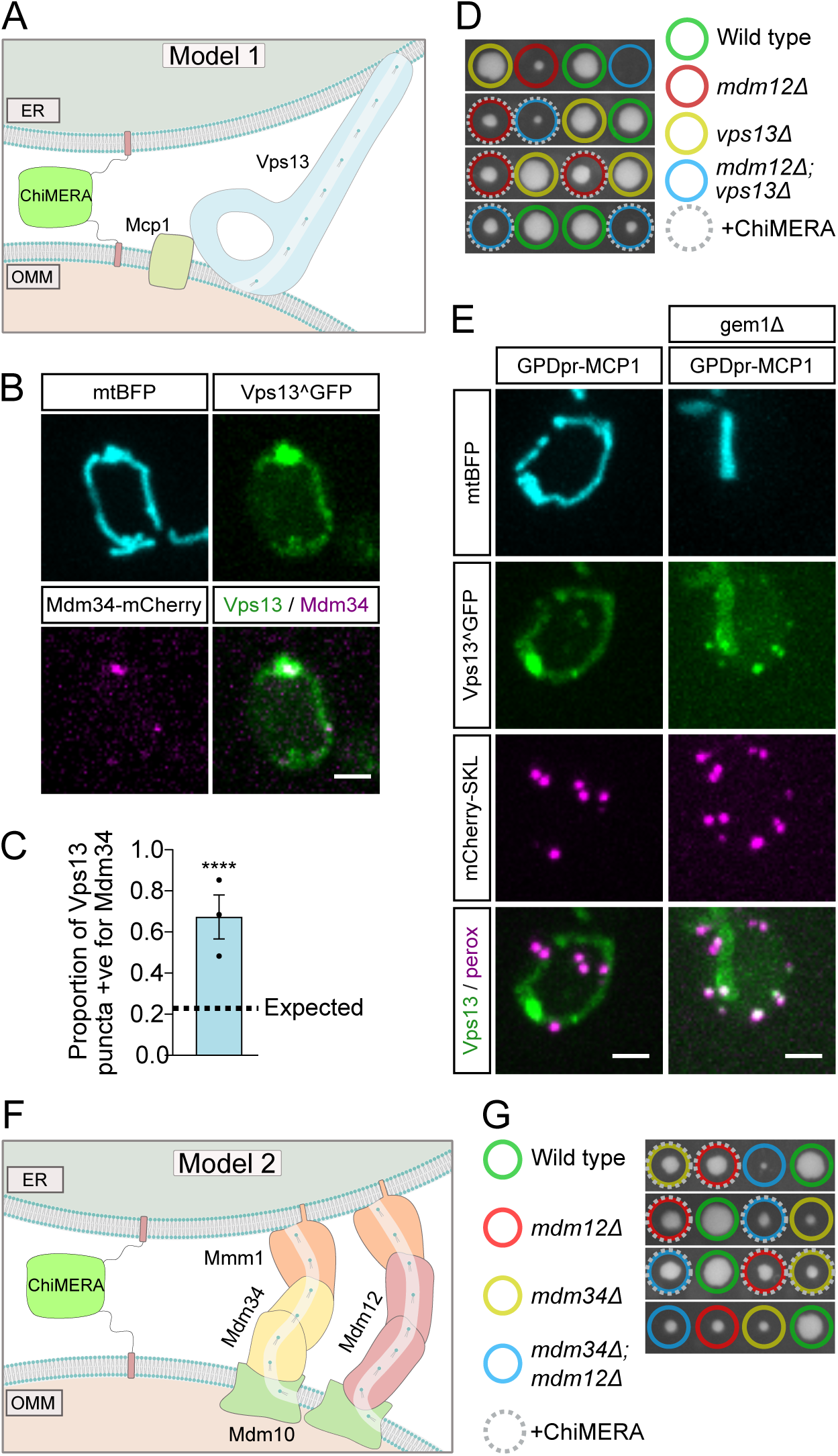
Vps13 and joint loss of Mdm12-Mdm34 are not required for ERMES rescue by ChiMERA. (A) Schematic showing the model in which Vps13-Mcp1 can transfer lipids at ChiMERA-induced ER-mitochondria contact sites. (B) Representative images of Vps13^GFP colocalizing with Mdm34-mCherry upon Mcp1 overexpression. (C) Quantification of the expected and observed extent of colocalization of Vps13^GFP with mdm34-mCherry. Statistical significance was quantified with a Chi-squared test. **** denotes p<0.0001. (D) Representative tetrads from the sporulation of *MDM12/mdm12*Δ *VPS13/vps13*Δ diploids expressing ChiMERA. (E) Representative images of Vps13^GFP subcellular localization in comparison to mitochondria (mtBFP) and peroxisomes (mCherry-SKL) in Mcp1 overexpression conditions, both with and without Gem1. (F) Schematic of Model 2, whereby Mdm12 and Mdm34 are functionally redundant upon ChiMERA-induced tethering of the ER to the mitochondria. (G) Representative tetrads from the sporulation of *MDM12/mdm12*Δ *MDM34/mdm34*Δ diploids expressing ChiMERA. Scale bars are 2 μm.

The localization of Vps13 to ER-mitochondria contact sites makes Vps13 a strong candidate for a lipid-transfer protein at these sites. If ChiMERA rescued ERMES deficiency through promoting lipid transfer by Vps13 at ER-mitochondria contact sites, ChiMERA expression would be expected to be ineffective in *vps13*Δ cells. To test this idea, we sporulated and tetrad-dissected an *MDM12/mdm12*Δ*; VPS13/vps13*Δ heterozygous diploid strain, transformed with a ChiMERA-expressing plasmid. As ERMES deficiency is synthetic lethal with *vps13*Δ (Lang et al., 2015), we expect that, if Vps13 was required for ChiMERA-mediated rescue, *mdm12*Δ*/vps13*Δ cells would be inviable irrespective of ChiMERA expression. While meiotic tetrad dissection showed that *mdm12/vps13* double knockout cells were not viable (Figure 2D), expression of ChiMERA led to a consistent growth restoration in these double knockout cells. Therefore, ChiMERA-induced rescue of ERMES deficiency does not require Vps13-dependent lipid transport between the ER and mitochondria, ruling out Model 1.

Instead of playing a role in the ERMES rescue pathway, the bright Vps13 foci observed upon Mcp1-overexpression appeared to be mitochondria-derived compartments (MDCs), mysterious outer-membrane-enriched organelle protrusions involved in quality control, longevity and resistance to amino acid stress (Schuler et al., 2021). Like MDCs, Vps13-foci formed at or in proximity of ERMES foci, and sometimes exhibited a doughnut shape (Supp. Figure 1D). Development of these foci was strictly dependent on Gem1, a calcium-binding GTPase (Frederick et al., 2004) and facultative components of the ERMES complex (Kornmann et al., 2011; Covill-Cooke et al., 2024; English et al., 2020) — Gem1 deletion abrogated the punctate localization of Vps13, resulting in a diffuse mitochondrial signal alongside some enrichment in foci which colocalized with peroxisomes (Figure 2E). Vps13 localization to peroxisome has been previously observed but its function is not understood (John Peter et al., 2017). We therefore conclude that Vps13’s localization to ER-mitochondrial contacts sites is more likely associated with MDC biology and unconnected to the ChiMERA-induced rescue of ERMES deficiency.

### Redundancy between Mdm12 and Mdm34 does not explain ChiMERA-mediated ERMES deficiency rescue

The expression of ChiMERA can rescue the growth defect observed in single deletion of either *mdm12* or *mdm34* (Kornmann et al., 2009). Model 2 implies that ER-mitochondria tethering by ERMES requires all four members of the complex, but that the functions of Mdm12 and Mdm34 in lipid transfer are redundant (Figure 2F), which becomes apparent when artificial tethering is provided by ChiMERA. Thus, one would expect that ChiMERA could rescue growth of *mdm12*Δ or *mdm34*Δ single mutants but not of *mdm12*Δ*/mdm34*Δ double mutants. To test this hypothesis, *MDM12/mdm12*Δ*; MDM34/mdm34*Δ heterozygous diploids were transformed with a ChiMERA-expressing plasmid and cell growth was examined following sporulation and tetrad dissection. ChiMERA expression elicited a similar rescue in *mdm12*Δ*/mdm34*Δ double knockout cells to that obtained in single deletion yeast (Figure 2G). We therefore conclude that Mdm12 and Mdm34 are not redundant for lipid transport, ruling out model 2.

### The SMP-domain of Mmm1 is sufficient for lipid transfer to mitochondria

Contrary to deficiencies in Mdm12, Mdm34 and combination thereof, deficiency in Mmm1 and Mdm10 are poorly, if at all, rescued by ChiMERA expression (Kornmann et al., 2009), indicating that these components play a special role beyond that of providing tethering force. We therefore hypothesised that Mmm1 and Mdm10 might be able to form a dimer that is functional in lipid transfer (Model 3, Figure 3A). In fact, AlphaFold3 predicts a highly confident Mmm1-Mdm10 heterodimer in an orientation compatible with lipid transfer between two membranes and of an appropriate size to fit within the narrow ER-mitochondria gap induced by ChiMERA-mediated tethering (Figure 3B). Since Mdm10 is devoid of a lipid transport domain, Mmm1 – which itself is ER-anchored - appears to be sufficient for interorganelle lipid transport, with Mdm10 serving as a recruitment factor on the mitochondrion.

**Figure 3:**
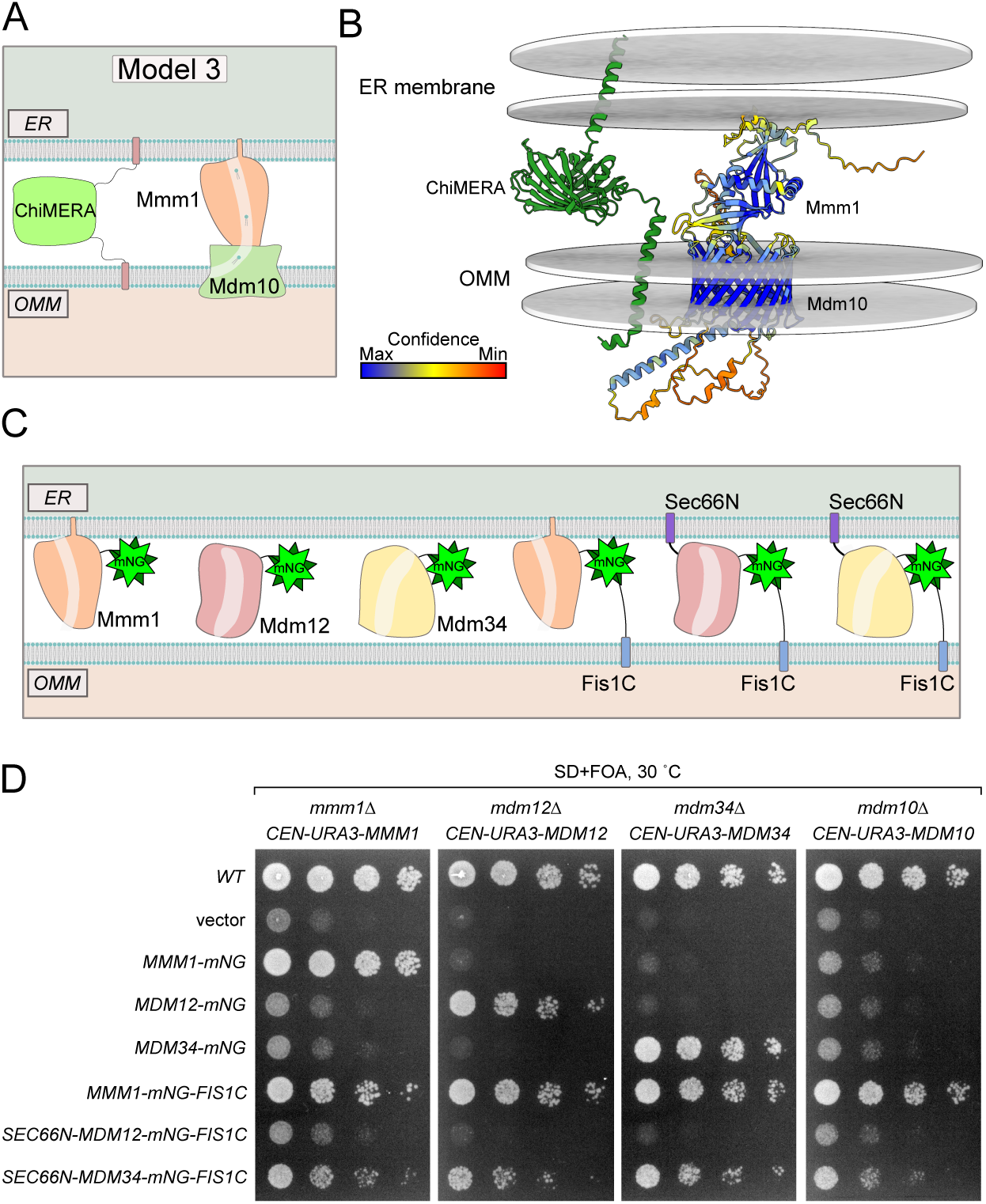
Artificial tethering of Mmm1 to mitochondria can rescue loss of ERMES. (A) Schematic of model 3 showing that a Mmm1-Mdm10 dimer is functional upon additional tethering. (B) AlphaFold3 prediction of Mmm1 and Mdm10 dimer. ChiMERA was added manually along with the ER and outer mitochondrial membrane (OMM). (C) Schematic of constructs used in (D). (D) Spot assay of single ERMES deletion yeast expressing artificially tethered ERMES members.

We thus asked whether Mmm1 artificially tethered to mitochondria could functionally replace the core subunits of the ERMES complex. To achieve this, we expressed Mmm1 fused to the fluorescent mNeonGreen (mNG) protein with a C-terminal OMM-targeting transmembrane domain (Fis1-TM, hereafter Mmm1-mNG-Fis1C) (Figure 3C). Interestingly, Mmm1-mNG-Fis1C could rescue not only the growth defects of a *mmm1*Δ strain, but of all the single ERMES deletion mutant strains including, unlike ChiMERA, *mdm10*Δ (Figure 3D), indicating that Mdm10 becomes dispensable when Mmm1’s SMP domain is artificially targeted to the OMM. We tested whether Mdm12 and Mdm34 could equally rescue ERMES deficiency when artificially targeted to ER-mitochondria contact sites. To do so, we fused both proteins to both an N-terminal ER-targeting transmembrane domain (Sec66-TM) and a C-terminal mNG with the OMM-targeting Fis1-TM (Figure 3C). Mdm34, targeted both mitochondria and the ER, could rescue the growth defects of *mmm1*Δ, *mdm12*Δ, and *mdm10*Δ strains, in addition to its cognate deletion strain, but to a lesser extent. Mdm12 failed to rescue the growth of even the *mdm12*Δ strain in the same setup (Figure 3D).

Since Mmm1-mNG-Fis1 could rescue each ERMES deficient strain individually, we next tested whether it could complement the combined quadruple ERMES deletion mutant (*mmm1*Δ*/mdm12*Δ*/mdm34*Δ*/mdm10*Δ). Indeed, Mmm1-mNG-Fis1C, could functionally complement the quadruple ERMES deletion mutant (Figure 4A). In addition to the growth defect, Mmm1-mNG-Fis1C also rescued the mitochondrial morphology phenotype observed upon ERMES deletion. Neither Mdm12 nor Mdm34 expressed in the same experimental setup rescued either growth or mitochondrial morphology (Figure 4B).

**Figure 4:**
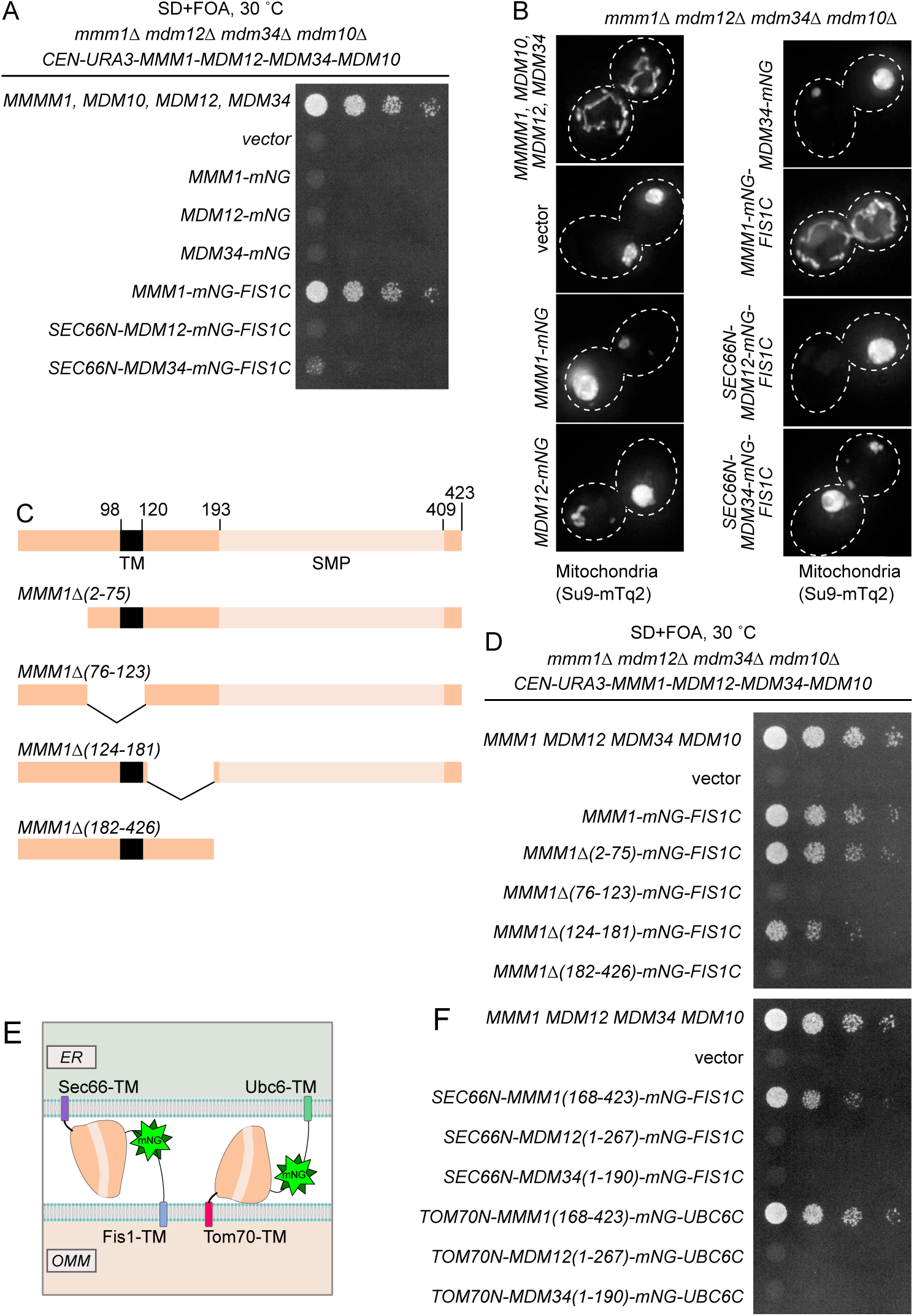
SMP domain of Mmm1 is sufficient for lipid transfer. (A) Spot assay of ERMES quadruple deletion yeast with and without the expression of artificially tethered ERMES members. (B) Representative images of mitochondria from the yeast in (A). Schematic of Mmm1 truncations constructs used in (D). (D) Spot assay of *mmm1*Δ*mdm12*Δ*mdm34*Δ*mdm10*Δ yeast expressing truncations of Mmm1. (E) Schematic of constructs used in (F). (F) Spot assay of *mmm1*Δ*mdm12*Δ*mdm34*Δ*mdm10*Δ yeast strains expressing tethering constructs with different transmembrane and SMP domains.

Mmm1 consists of an N-terminal ER-luminal domain dispensable for function (residues 1-97), an ER-inserted transmembrane domain (residues 98-120), and a cytosolic SMP domain (193-409), connected by linker sequences. To identify which of these domains are essential for the observed rescue activity, we generated a series of partial truncation and fusion constructs (Figure 4C). Deletion of the ER-luminal region (2-75) or the cytosolic linker region (124-181) did not affect the rescue activity, consistent with these domains being dispensable for Mmm1 function, yet deletion of the TM domain (76-123) or the SMP domain (182-426) abolished the rescuing ability of the construct (Figure 4D). To assess whether Mmm1’s TM domain played a special role beside targeting, we replaced it with Sec66-TM (Figure 4E) expressed from the ADH1 promoter. This substitution did not compromise the rescue activity (Figure 4F), indicating that Mmm1 TM domain does not serve a special function besides ER attachment. To assess whether the order of the targeting sequences on the protein was important, we designed constructs where the N-terminal side of Mmm1’s SMP domain was attached to an OMM-targeting domain (Tom70-TM) and its C-terminus to an ER-targeting domain (Ubc6) (Figure 4E). Even in that inverted configuration, Mmm1’s SMP domain could rescue the mutant growth defects (Figure 4F). By contrast, the SMP domain of Mdm12 or Mdm34 did not rescue the growth defect of the mutant yeast in either configuration (Figure 4F). To conclude, the SMP domain of Mmm1 can function alone in lipid transfer between the ER and mitochondria, outside of the ERMES complex, and Mdm10 is solely responsible for tethering the complex, either full or partial, to the mitochondrion.

## Discussion

The ERMES complex was originally discovered as an ER-mitochondria tether because its function could be substituted by an artificial ER-mitochondria tethering protein – ChiMERA (Kornmann et al., 2009). Further structural and functional studies have unambiguously demonstrated that ERMES catalyses interorganelle lipid transport via lipid transport domains in Mmm1, Mdm12 and Mdm34 (Kawano et al., 2018; Jeong et al., 2017; AhYoung et al., 2015, 2017; John Peter et al., 2022). This created a puzzling conundrum; how could an artificial tether without lipid transport activity rescue a compositionally rigid protein complex with demonstrated lipid transport activity? Our data rule out a redundant role of Mdm12 and Mdm34 in lipid transport and an involvement of Vps13 in ChiMERA-mediated rescue of ERMES deficiency. Vps13 involvement in ChiMERA-mediated ERMES suppression was a tempting model, since bolstering the Mcp1/Vps13 pathway (either by point mutations in Vps13 or by Mcp1 overexpression) is the only other understood means to achieve ERMES rescue (Park et al., 2016; Lang et al., 2015; Kojima et al., 2016; Tan et al., 2013). Moreover,Vps13 was shown to harbour lipid transport activity (Li et al., 2020; Adlakha et al., 2022; Kumar et al., 2018).

Vps13 localizes to foci at ER-mitochondria contacts in a Mcp1-overexpression- and Gem1-dependent fashion. Recently, ERMES foci have been shown to be sites of budding of MDCs — protrusions of OMM enriched in various OMM proteins that are important to survive amino acid stresses (Schuler et al., 2021). ER-mitochondrial Vps13 foci are most likely MDCs for three reasons: 1) they colocalized with Mdm34-mCherry-labelled ERMES foci, 2) they were more prominent and adopted a doughnut-like appearance when cells reached the saturation phase and 3) their appearance was entirely dependent on Gem1. In fact, a microscopy-based screen identified Mcp1 as being enriched at MDCs (Hughes et al., 2016). That Vps13-mediated lipid transport plays a role in MDC biology seems plausible, given the importance of lipid homeostasis on MDC formation (Xiao et al., 2024).

We here argue that Mmm1 and Mdm10 might, by themselves suffice to provide a core lipid exchange platform, upon additional tethering force (Model 3; Figure 1D). We find that the SMP domain of Mmm1 alone can rescue the complete loss of all ERMES members providing it is recruited to ER-mitochondria contact sites, by fusion to ER and mitochondria targeting motifs at its N and C termini. This Model 3 could explain why, contrary to deficiencies in Mdm12, Mdm34 and combination thereof, deficiency in Mmm1 and Mdm10 are poorly, if at all, rescued by ChiMERA expression, indicating that these components play a special role that cannot be substituted by tethering force. This two-protein model would also explain the results of a screen in fission yeast that found that Mmm1 overexpression could rescue the lethality of cells defective for Mdm12, Mdm34, and double mutants thereof, but not for Mdm10 (Li et al., 2019).

The functionality of the subcomplexes we observe, in particular of a minimal Mmm1-Mdm10 complex, imply that the ERMES heterotetramer observed by cryotomography (Wozny et al., 2023) might not be the only functional unit formed by ERMES proteins. It is possible that ERMES consists of a family of complexes ranging from the proposed Mmm1-Mdm12-Mdm34-Mdm10 unit to a minimal Mmm1-Mdm10 unit, which is not *per se* sufficient for function but can be made sufficient by the addition of extra tethering force, either via ChiMERA expression in *mdm12/mdm34* mutant cells, or by the presence of additional ‘complete’ ERMES complexes in wild type cells. The composition, subcellular localization, lipid specificity and overall contribution in lipid exchange of the Mmm1-Mdm10 subcomplex will need to be clarified. It might, for instance, be able to span different spacings as found at ER-mitochondria interfaces. Finally, this model provides an evolutionary path from a simple two-component lipid transport complex to a more complex four-subunit one.

## Materials and Methods

### Plasmids and yeast strains

**Table 1:**
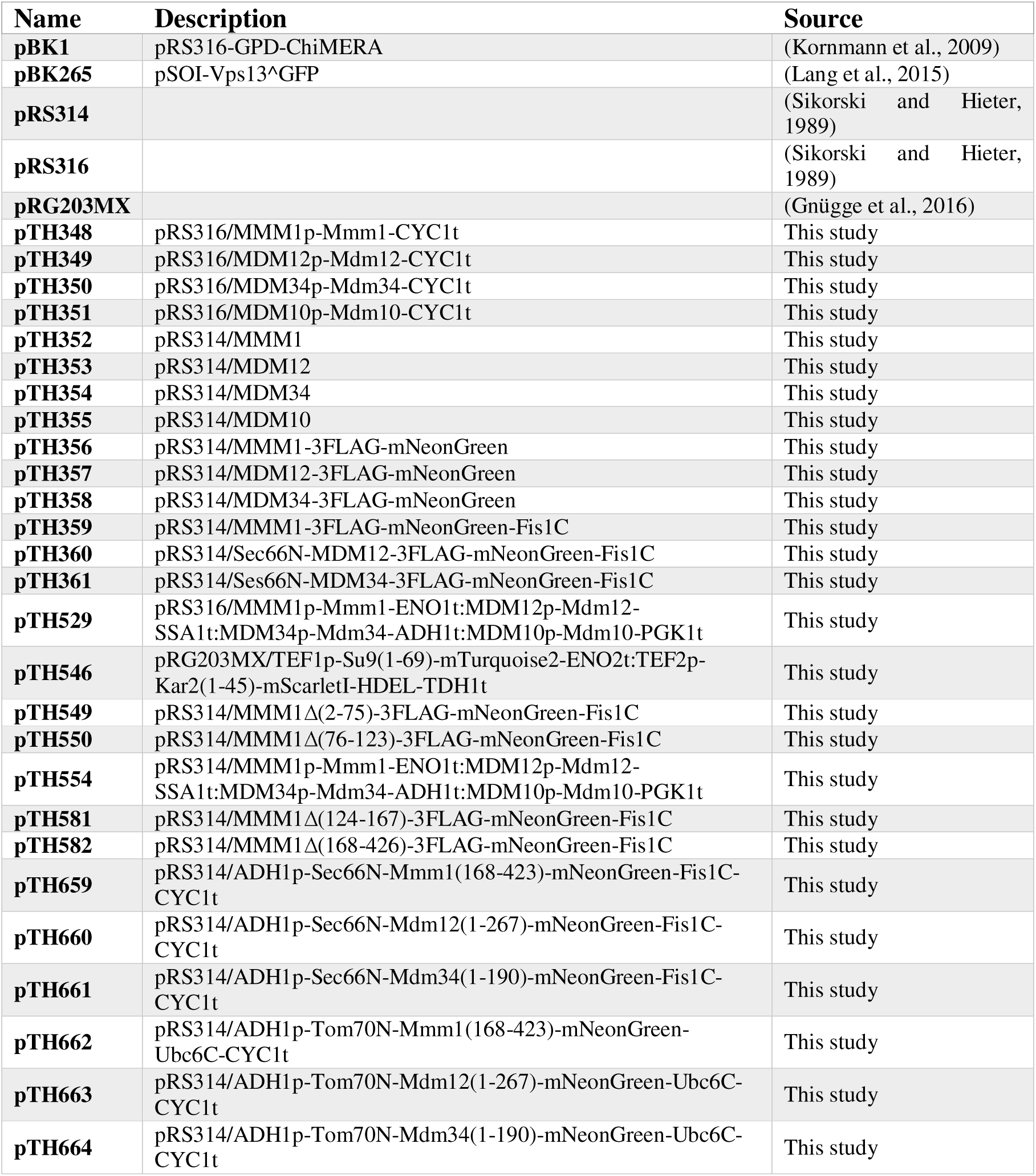
All plasmids used in this study.

Yeast gene deletions with selective markers were achieved as previously described (Longtine et al., 1998). Overexpression of the endogenous *MCP1, VPS39* and *MMM1* genes was achieved by integrating a GPD promoter cassette as previously described (Janke et al., 2004).

Marker-free gene deletions were achieved by the CRISPR/Cas9 system as previously described (Soreanu et al., 2018).

**Table 2:**
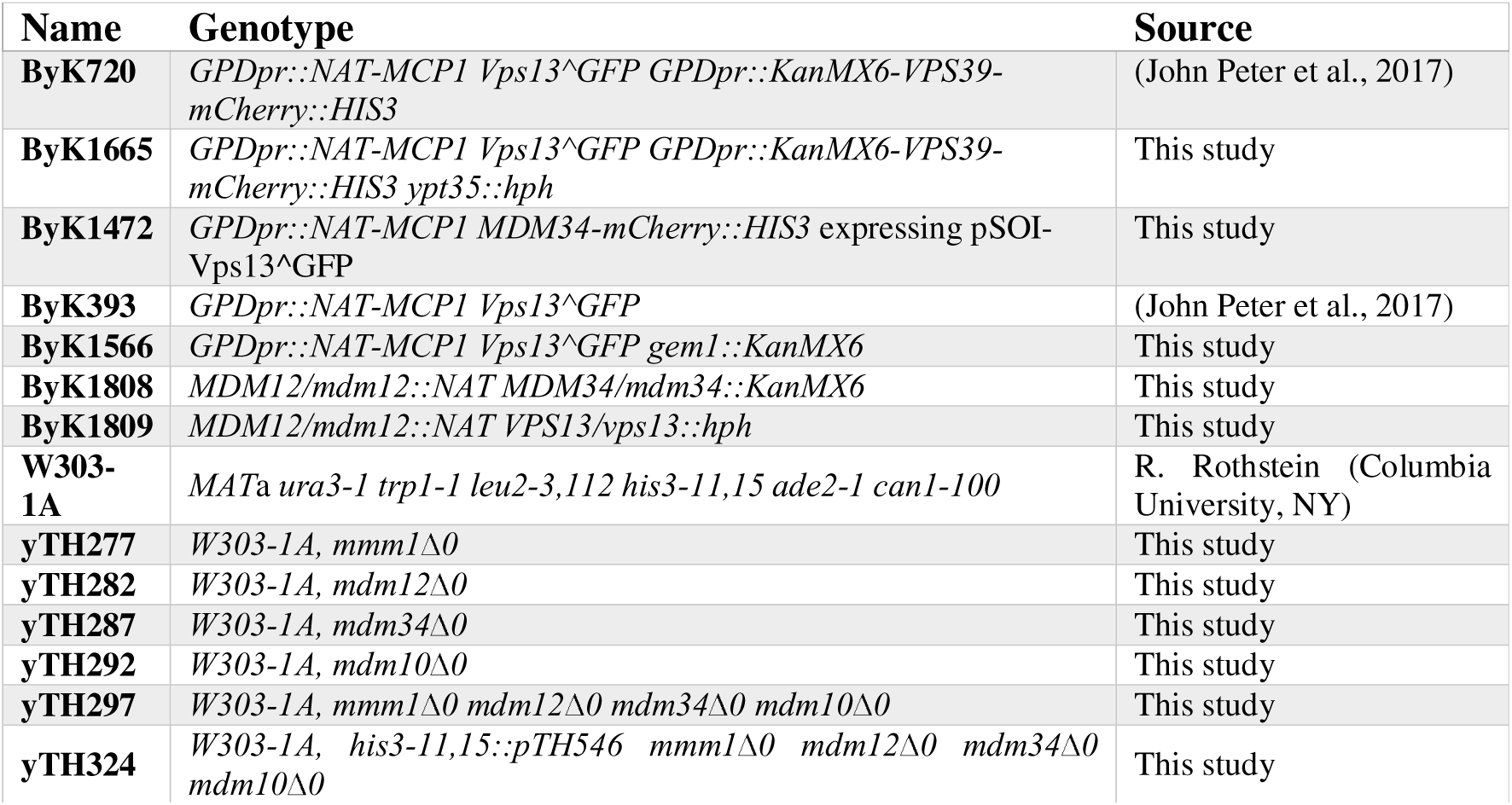
Yeast strains used in this study.

### Media

Yeast cells were cultured in SD media (0.67% yeast nitrogen base without amino acids, 2% glucose) supplemented with 20 µg mL^−1^ each of adenine sulfate, L-histidine-HCl, L-tryptophan, and uracil and 30 µg mL^−1^ each of L-leucine and L-lysine-HCl. For spot assay, 1 mg mL^−1^ 5-fluoroorotic acid was added. For microscopic observation, the concentration of adenine sulfate was doubled.

### Spot assay (plasmid shuffling)

Yeast strains were grown to saturation in SD lacking tryptophan and uracil, then diluted 100-fold with SD lacking tryptophan, and cultured for 24 h. Then, the cultures were serially diluted 5-fold, spotted onto agar plates containing 5-fluoroorotic acid, and incubated at 30°C for 48 h.

### Tetrad analysis

Diploid yeast strains were sporulated by patching cells onto agar plates containing 1% yeast extract, 2% peptone and 1% potassium acetate. After appearance of tetrads (5-7 days), they were dissected onto YPD and subsequently genotyped by replica plating onto media to select for the markers of interest.

### Fluorescence microscopy

For Figure 2: Saturated cultures of yeast were reseeded in appropriate selective media at a very low density so that they would still be in log phase after at least 16 hours of growth. Then, approximately 500,000 cells were plated on a microscope slide with a coverslip on top. Images were obtained using an UltraView IX81 Olympus spinning disc confocal microscope equipped with a 100x oil immersion objective lens (NA=1.4). Image acquisition, using Volocity software, was carried out at room temperature. For Figure 3: Yeast strains were grown to log phase and placed on a microscope slide with a coverslip. Cells were observed at room temperature using a DeltaVision Elite system (Cytiva) equipped with a 100× objective lens (UPLSAPO, NA/1.40; Olympus) and a sCMOS camera (Edge5.5; PCO). Z-sections were captured every 0.2 or 0.4 μm from the top to the bottom surface of yeast cells. Deconvolution was performed using SoftWoRx software (Cytiva), and the acquired images were processed and analyzed with Fiji software. For Supplementary Figure 1D: Images were acquired using a DeltaVision MPX microscope (Applied Precision) equipped with a 100× 1.40 NA oil UplanS-Apo objective lens (Olympus), a multicolor illumination light source, and a CoolSNAPHQ2 camera (Roper Scientific). Image acquisition was done at RT. Images were deconvolved with SoftWoRx software using the manufacturer’s parameters.

### Image analysis and statistics

To ascertain whether Vps13^GFP puncta preferentially localize at ERMES foci, first, cells were scored whether they had colocalization of Vps13^GFP and Mdm34-mCherry signal. This was determined to be the case if at least half of the Mdm34-mCherry pixels overlapped with the Vps13^GFP puncta signal. The percentage of cells with colocalization for each repeat was then compared to the expected probability that a Vps13^GFP puncta would randomly overlap with Mdm34-mCherry. This expected value was calculated by quantifying the size of the thresholded Vps13^GFP puncta and dividing it by the mitochondrial mass (calculated from the thresholded signal of the mitochondrial marker mtBFP), i.e., the chance that a Vps13^GFP puncta would fall at any position of the mitochondria. The percentage of cells showing colocalization (observed) was then compared to this expected value using a Chi-squared test.

### Structural predictions

Structural prediction of the Mmm1-Mdm10 dimer were made using AlphaFold3 (Abramson et al., 2024) and visualised using ChimeraX (Pettersen et al., 2021).

## Acknowledgments

The authors gratefully acknowledge the Micron Advanced Bioimaging Facility (supported by Wellcome Strategic Awards 091911/B/10/Z and 107457/Z/15/Z) for their support & assistance in this work. This work was funded by Wellcome Trust grant 214291/Z/18/Z (awarded to BK), JSPS KAKENHI 15H05705 and 2222703 (to T.E.), JSPS KAKENHI 17K18230 and 25840020 (to S.K.), JST CREST grant JPMJCR12M1 (to T.E.), AMED CREST grant 21gm1410002h0002 (to T.E.), and a grant from Takeda Science Foundation (to T.E.). T.H. (23KJ2067) was supported by a Research Fellowship for Young Scientists from the Japan Society of the Promotion of Society.

## Supplementary figure legends

**Supplementary Figure 1: Vps13 forms puncta on mitochondria.** (A) Comparison of transposon density maps for *MCP1* and *YPT35* in wild type, *VPS13^D716H^* and *mmm1*Δ *VPS13^D716H^* yeast using SATAY. (B) Representative images of Vps13^GFP localization to vCLAMPs in Mcp1 overexpression conditions, both with and without Ypt35. mtBFP is a mitochondrial marker. (C) Representative images of Vps13^GFP both with and without the overexpression of *MCP1*. Scale bars are 2 μm. (D) Individual Z planes of Vps13^GFP when cells are in stationary phase. Scale bars are 5 μm.

