## Supplementary figures and images for "Compositional Flexibility of the ER-Mitochondria Encounter Structure"

### Supp. Figure 1

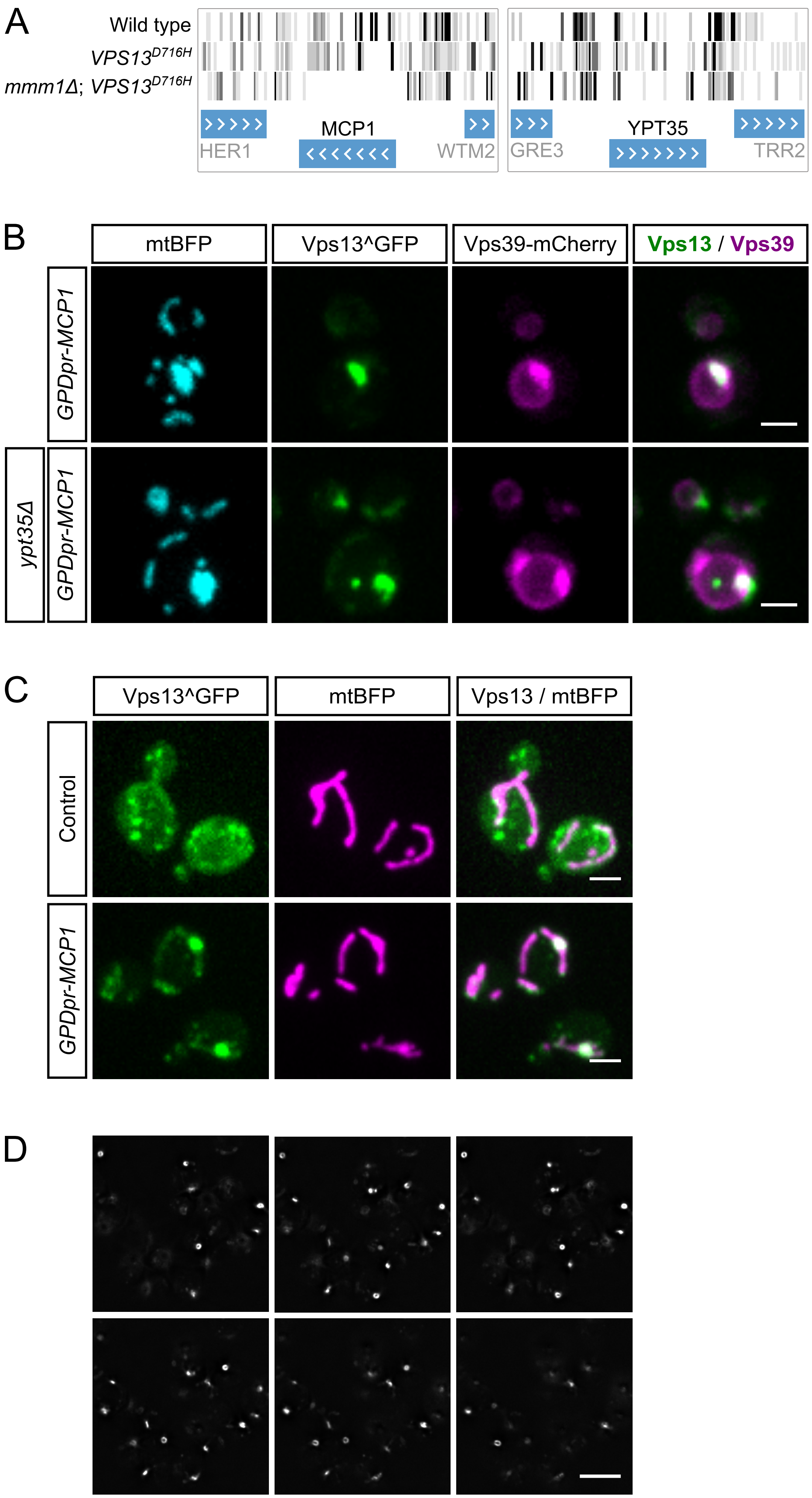
